# Quantifying the Impact of Co-Housing on Murine Aging Studies

**DOI:** 10.1101/2024.08.06.606373

**Authors:** Alison Luciano, Gary A. Churchill

**Affiliations:** The Jackson Laboratory, Bar Harbor, ME, USA

**Keywords:** aging, study design, co-housing

## Abstract

Analysis of preclinical lifespan studies often assume that outcome data from co-housed animals are independent. In practice, treatments, such as controlled feeding or putative life-extending compounds, are applied to whole housing units, and as a result the outcomes are potentially correlated within housing units. We consider intra-class (here, intra-cage) correlation in three published and two unpublished lifespan studies of aged mice encompassing more than 20 thousand observations. We show that the independence assumption underlying common analytic techniques does not hold in these data, particularly for traits associated with frailty. We describe and demonstrate various analytical tools available to accommodate this study design and highlight a limitation of standard variance components models (i.e., linear mixed models) which are the usual statistical tool for handling correlated errors. Through simulations, we examine the statistical biases resulting from intra-cage correlations with similar magnitudes as observed in these case studies and discuss implications for power and reproducibility.

## Introduction

Lifespan is a complex trait that is influenced by multiple genetic and environmental factors. Understanding the determinants of lifespan and aging is a major challenge in biomedical research, with implications for human health and longevity. However, the reproducibility and generalizability of lifespan studies are limited by the variability and heterogeneity of experimental conditions, the choice of animal model, the design of the study, the measurement of lifespan, and the reporting of data.

One of the most widely used animal models for lifespan research is the mouse, which offers several advantages, such as accelerated study timelines and availability of genetic tools. However, murine lifespan studies also face several challenges, including the potential confounding effects of housing conditions on outcomes. Commonly used statistical methods for evaluating treatment effects in murine lifespan data assume that the survival of each individual is independent of the survival of others. This assumption may not hold, as mice are social animals that interact with each other and may influence each other’s lifespan through behavioral, physiological, or immunological mechanisms.

In intervention studies, treatments are often administered in food or water and, as such, the treatments are being applied to housing units and not to individual animals [1]. This is a common experimental design known as a cluster-randomized trial (CRT). Analysis methods for CRTs that account for within-cluster correlations have not been widely adopted in pre-clinical lifespan studies [2]. Appropriate methods may include linear mixed models [3, 4], generalized estimating equations [5, 6], additive hazards mixed models [7], sandwich variance estimators [8], semiparametric mixed-effects models [9, 10], parametric mixed-effects models [11], and copula models [12]. Each approach has its own assumptions, advantages, and disadvantages, depending on the research question, the study design, and the data characteristics. Nonetheless, it has been a common practice to analyze murine lifespan studies as if the individual mice were randomized to treatments, ignoring the possible consequences of intra-class correlation (ICC). This has implications for power and reproducibility of murine aging studies.

Our aim in this work is to determine whether the lifespans and longitudinally collected phenotypes of co-housed animals are independent and to better understand the implications of co-housing for study design and analysis outcomes (**Fig. 1**). We performed a secondary analysis of mouse lifespan data from six large cohorts. The assembled database comprises over 20,000 observations with lifespan and cage assignment data. These studies include mice of both sexes representing 42 inbred strains and two outbred populations. The studies were carried out in different facilities, at different times and with a variety of housing configurations. We carried out simulations to evaluate analysis methods, to estimate samples size, and to assist researchers in obtaining unbiased results by providing recommendations for design and analysis of group-randomized preclinical lifespan trials.

**Figure 1:**
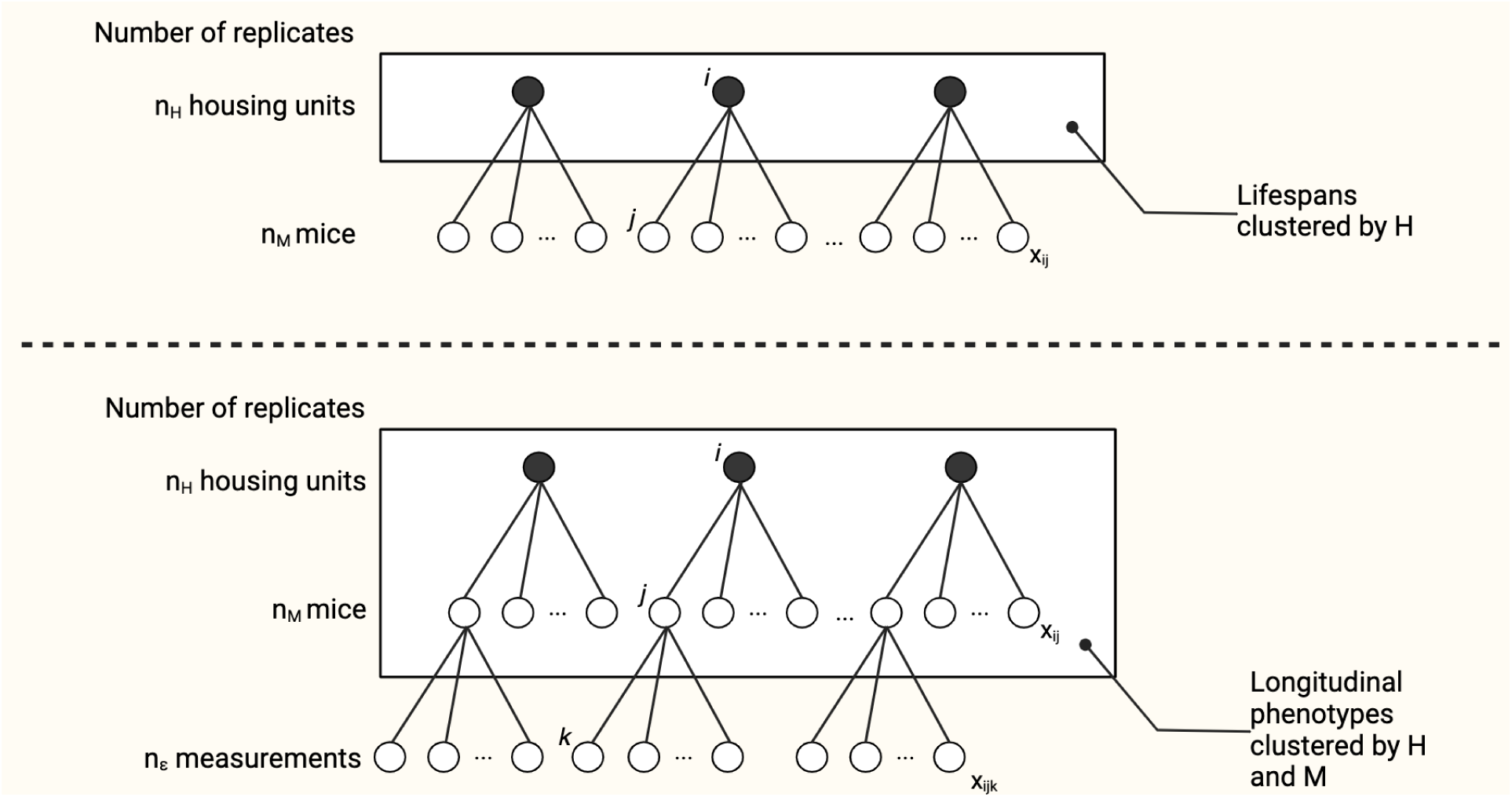
Clustering of lifespan and phenotypic trajectories in murine aging studies. Analysis of preclinical lifespan studies often assumes that outcome data from co-housed animals are independent. In practice, treatments, such as controlled feeding or putative life-extending compounds, are applied to whole housing units, and as a result the outcomes are potentially correlated either within housing units H (as in lifespans) or within H and mouse M (as in longitudinal phenotypes). We quantify intra-class correlation in a pooled database of murine aging studies encompassing more than 20 thousand observations.

## Results

### A murine lifespan database

Murine lifespan studies are a cornerstone of research in the basic biology of aging. Our objective was to determine if survival of co-housed individuals is independent and to explore the implications of ICC. We obtained data from four published and two unpublished murine lifespan cohorts for which housing identifiers were available totaling 22,385 mice (**Table 1**). These studies include mice of both sexes representing 42 inbred strains and two outbred populations. The studies were carried out in different facilities, at different times and with a variety of housing configurations. The mice included in the database were genetically diverse, including 3,763 inbred mice (JAX32+JAXCC studies) and 18,622 outbred mice (JAXDO+DRiDO+UTITP+JAXITP). Among the outbred mice, we obtained data for 17,062 UM-HET3 mice (UTITP+JAXITP) and 1,560 DO mice (JAXDO+DRiDO). Both sexes are well represented (50.8% female). With these data we were able to evaluate independence of murine lifespan across more that 20,000 mice and nearly 6,000 housing units. Survival outcome varied by study, site, randomization group, strain, and sex (**Fig. 2**).

**Figure 2:**
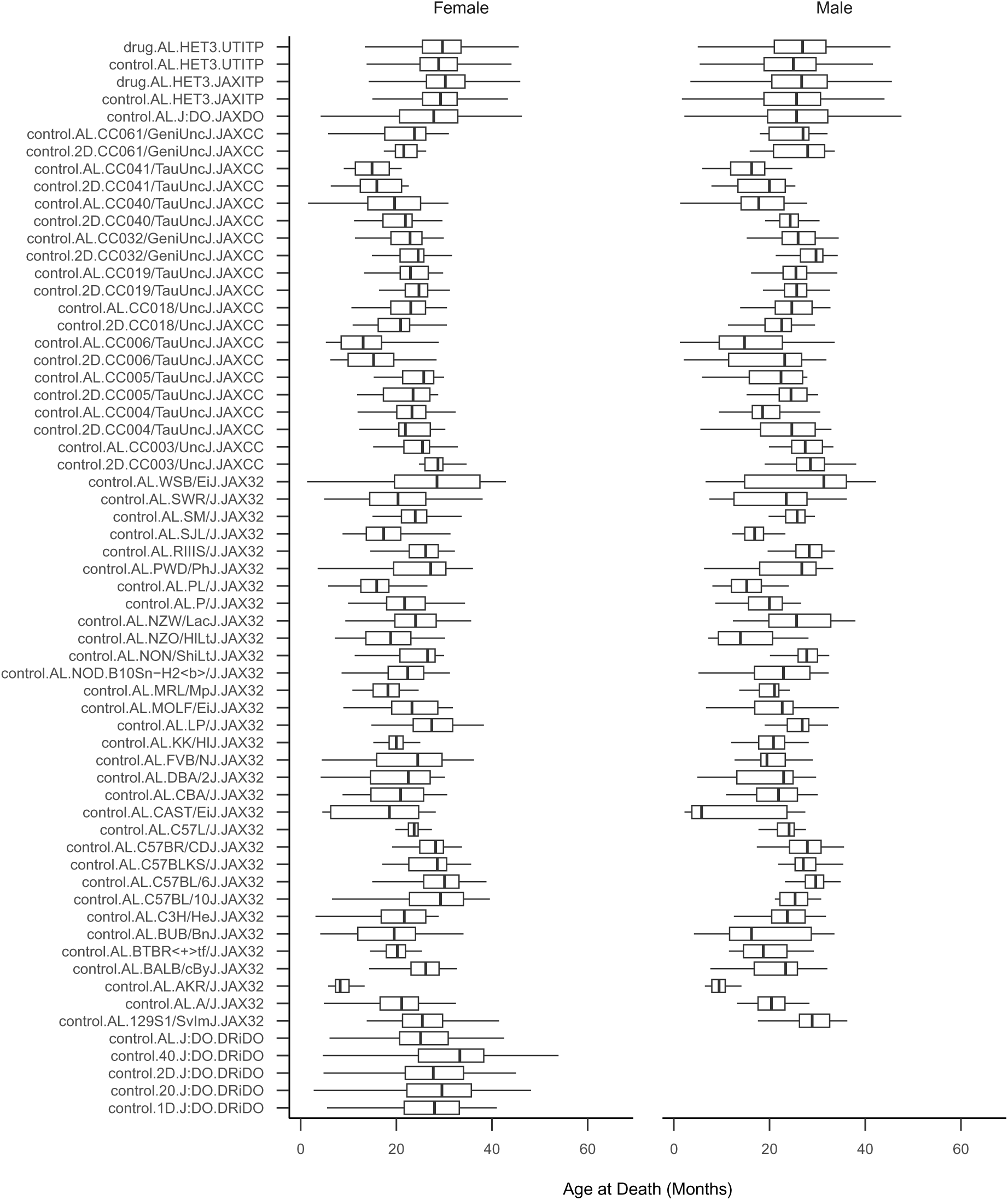
Survival by study, site, randomization group, strain, and sex. Boxplots show the primary outcome, lifespan, across study, randomization group, strain, and sex; N strata = 120. Lifespans shown are among mice that died of natural causes. Strata are defined as follows: [drug].[diet].[strain].[primary study]. ‘drug’ indicates treatment condition in ITP study as either control or drug, where control = assigned to no treatment and drug = assigned to a treatment (treatments combined, regardless of efficacy); ‘diet’ indicates one of either AL (ad libitum diet), 2D (2-day intermittent fast), 1D (1-day intermittent fast), 20 (20% calorie restriction), or 40 (40% calorie restriction); strains are as written; primary study is one of either UTITP (University of Texas Health San Antonio Intervention Testing Program), JAXITP (The Jackson Laboratory Intervention Testing Program), JAXCC (The Jackson Laboratory Collaborative Cross Longitudinal Study), JAXDO (The Jackson Laboratory Diversity Outbred Longitudinal Study), JAX32 (The Jackson Laboratory Inbred Strain Survey Study), or DRiDO (Dietary Restriction in Diversity Outbred Mice Study).

**Table 1:**
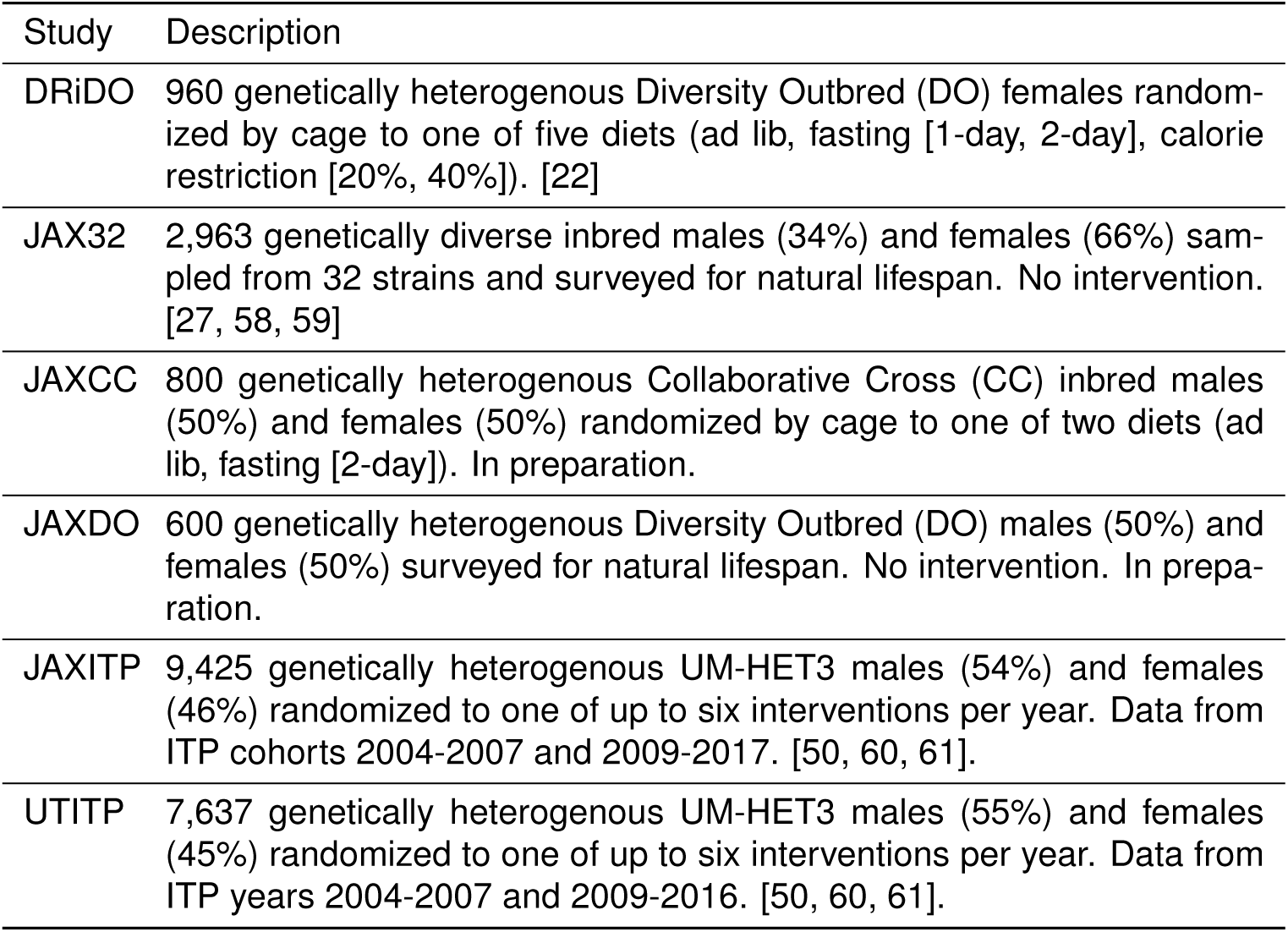
Study characteristics for the large database amassed for analysis, which includes lifespan and housing identification data for *>*20k aged mice. Database combined N = 22,385. Main outcome was age at natural death (lifespan) for all studies. Primary data were provided by the JAX and UT Nathan Shock Centers of Excellence in the Basic Biology of Aging. Study abbreviations: UTITP (University of Texas Health San Antonio Intervention Testing Program), JAXITP (The Jackson Laboratory Intervention Testing Program), JAXCC (The Jackson Laboratory Collaborative Cross Longitudinal Study), JAXDO (The Jackson Laboratory Diversity Outbred Longitudinal Study), JAX32 (The Jackson Laboratory Inbred Strain Survey Study), DRiDO (Dietary Restriction in Diversity Outbred Mice Study).

Housing densities varied by study. Mice were most commonly housed four per cage (45.4% of mice), then three per cage (36.0% of mice). All mice in the JAXDO study were housed 5 per cage and all mice in the DRiDO study were housed 8 per cage. Housing densities varied within other studies — the JAXITP study housing density was the most variable with a standard deviation of 1.25 mice (versus SD=0.933 [JAX32], SD=0.781 [JAXCC], and SD=0.487 [UTITP]). Housing density affects several factors associated with lifespan [13–17]. Mice are sometimes reassigned to new housing units during a study, often to mitigate aggressive behavior [18]. We have analyzed these data “as randomized” based on the original housing assignments when possible.

In the absence of confounders, we might expect that survival outcome would be largely explained by study factors, especially among inbred mice where genetic heritability of lifespan is accounted for. Yet after adjusting for study-specific covariates, we found consistently large dispersion among survival times within cage (median intra-cage IQR of residual survival (JAX32 [3.9 months]*<*JAXCC [4.6 months]*<*UTITP [5.5 months]*<*JAXITP [5.9 months]*<*JAXDO [8.8 months]*<*DRiDO [9.8 months]). As an illustration, see per-housing-unit adjusted lifespan distribution for JAXDO (**Supplementary Fig. 1**). In the figure, each boxplot represents one housing unit. Boxplots of lifespan by housing ID were sorted within study by median adjusted lifespan. Taller boxplots within study indicate cages with larger intra-cage variability and non-zero slope of the medians indicates inter-cage variability within study. While all studies will exhibit some inter-cage variability, those with greater intra-cage correlation will exhibit more pronounced slope of median residual lifespan.

### Lifespans of co-housed animals are not independent

The intraclass correlation (ICC; [19]) provides a quantified measure of the degree of clustering in lifespans by housing units. We considered two approaches to estimating ICC, linear mixed models (LMM) and generalized estimating equations (GEE). LMM assumes that co-housed mice share a random contribution to lifespan that is common to all mice within a housing unit and that varies between housing units. GEE explicitly models the covariance matrix based on group structure. Consequently, GEE is an unbiased estimator of ICC in that it can estimate positive, null, or negative ICC values while LMM is a biased ICC estimator in that it can estimate only positive or null ICC values, a distinction with important downstream effects on analytic results [20]. In the presence of negative ICC, LMM will constrain random effect variance to zero, thereby forcing the statistical model to assume data are independent.

We applied LMM and GEE methods to each of the studies to estimate ICC in lifespan outcome (**Table 2**, **Fig. 3**). GEE estimated a small negative intra-cage correlation in DRiDO data which was missed by the LMM estimator (and failed to converge using default settings). LMM results indicated positive intra-cage correlation in JAXCC data, while GEE results indicated these data were consistent with positive, null, or negative intra-cage correlation. LMM results indicated large positive intra-cage correlation in both JAXITP and UTITP relative to other studies; qualitatively similar results were obtained via GEE estimation. LMM results in JAX32 indicated relatively large positive intra-cage correlation and qualitatively similar results were obtained via GEE estimation. LMM results in JAXDO indicated these data were consistent with positive or (nearly) null intra-cage correlation while GEE results indicated these data were consistent with positive, null, or small negative intra-cage correlation. LMM estimation of ICC for the pooled database after adjustment for study indicated that overall lifespan data were weakly positively correlated within cage (ICC [95% CI]: 0.049 [0.038, 0.059]). Similarly specified GEE modeling also found ICC hovered around 0.05 (ICC [95% CI]: 0.045 [0.033, 0.057]). The two models approached equal precision (difference in ICC CI width consistently less than 0.1).

**Figure 3:**
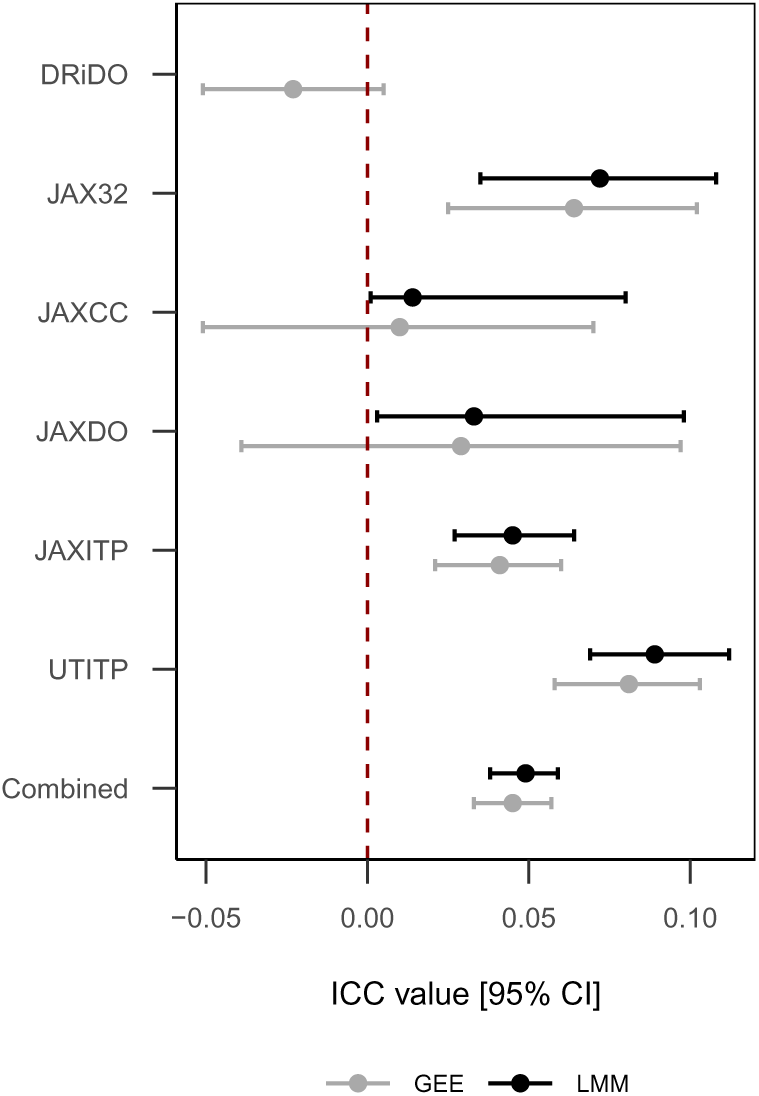
Intra-cage correlations by study and estimation method. LMM specifications: For DRiDO study, diet fixed effect and HID random effect (did not converge); survival in months adjusted for generation batch effect prior to modeling. For JAX32 study, sex fixed effect, strain and housing ID random effect. For JAXCC, Sex fixed effect, strain and housing ID random effects; survival in months adjusted for generation and cohort prior to modeling. For JAXITP and UTITP studies, sex and binary treatment/control fixed effects and housing ID random effect; survival in months adjusted for cohort prior to modeling. GEE specifications: Same as above for DRiDO, JAXDO, JAXITP and UTITP. In GEE analyses, JAX32 and JAXCC data were stratified by strain and the strain random effect term was dropped from the model to isolate the clustering effect of housing ID.

**Table 2:** Conditional (LMM) and marginal (GEE) estimation of intraclass correlations in lifespan outcome. The intraclass correlation coefficient (ICC) provides a quantified measure of the degree of clustering in lifespans by housing unit via linear mixed models (LMM) and generalized estimating equations (GEE). LMM conditions on random effects, while GEE integrates out random effects. Historical estimates of ICC can be used to inform sample size calculations for future CRTs.

| Study | LMM ICC (95% CI) | GEE ICC (95% CI) |
| --- | --- | --- |
| DRiDO | NA | -0.023 (-0.051, 0.005) |
| JAX32 | 0.072 (0.035, 0.108) | 0.064 (0.025, 0.102) |
| JAXCC | 0.014 (0.001, 0.08) | 0.01 (-0.051, 0.07) |
| JAXDO | 0.033 (0.003, 0.098) | 0.029 (-0.039, 0.097) |
| JAXITP | 0.045 (0.027, 0.064) | 0.041 (0.021, 0.06) |
| UTITP | 0.089 (0.069, 0.112) | 0.081 (0.058, 0.103) |
| Combined | 0.049 (0.038, 0.059) | 0.045 (0.033, 0.057) |

We used permutation testing to quantify significance of the observed ICC statistic. As recommended by R. A. Fisher [21], we treated the experimental unit assigned to treatments through randomization as the units that are permuted to assess ICC significance. In the experiments in the database, these units correspond to housing identifiers, as cages, not individual mice, are directly assigned to specific diets, sexes, or strains. Non-parametric statistical methods estimate significance levels of non-independence by cage that were not significantly different from the permuted (null) ICC distribution for DRiDO (p=0.887), JAXCC (p=0.401), and JAXDO (p=0.161), but were significantly different from the permuted ICC distribution (p*<*0.001) for JAX32, JAXITP, and UTITP (**Supplementary Fig. 2**). These results qualitatively correspond to GEE estimates of ICC significance.

### Longitudinally collected phenotypic trajectories of co-housed animals are not independent

Biomarkers are important indicators of normal and pathogenic aging biology and/or anti-aging therapeutic response. The DRiDO study, a comprehensive investigation of dietary restriction interventions in Diversity Outbred mice [22], offered an exceptional opportunity to explore clustering effects in aging biomarkers. Longitudinal phenotyping was conducted across multiple physiological domains, including weekly body weights, assessments of frailty index, grip strength, and body temperature every 6 months, and yearly evaluations such as metabolic cage analysis, body composition, echocardiogram, wheel running, rotarod performance, acoustic startle response, bladder function, fasting glucose levels, immune cell profiling, and whole blood analysis.

We applied LMM methods to all longitudinally collected phenotypic trajectories in DRiDO to quantify ICC across hundreds of physiologic measurements of aging (**Supplementary Table 1**). Examining maximum estimated ICC by physiological domain indicated positive intra-cage correlation could exceed 0.05 in several physiological domains, including frailty, metabolic cage, whole blood analysis, and wheel running. Domains with the top five median ICC included frailty, bladder function, metabolic cage, glucose, and wheel running (ICC=0.0674, 0.0356, 0.0302, 0.0227, and 0.0142 respectively), indicating these domains were sensitive to co-housing relative to other physiological domains (**Fig. 4**). Alopecia and whisker trimming may be induced by barbering among co-housed mice, providing a potential explanation for why these frailty indicators exhibited more pronounced effects than other traits.

**Figure 4:**
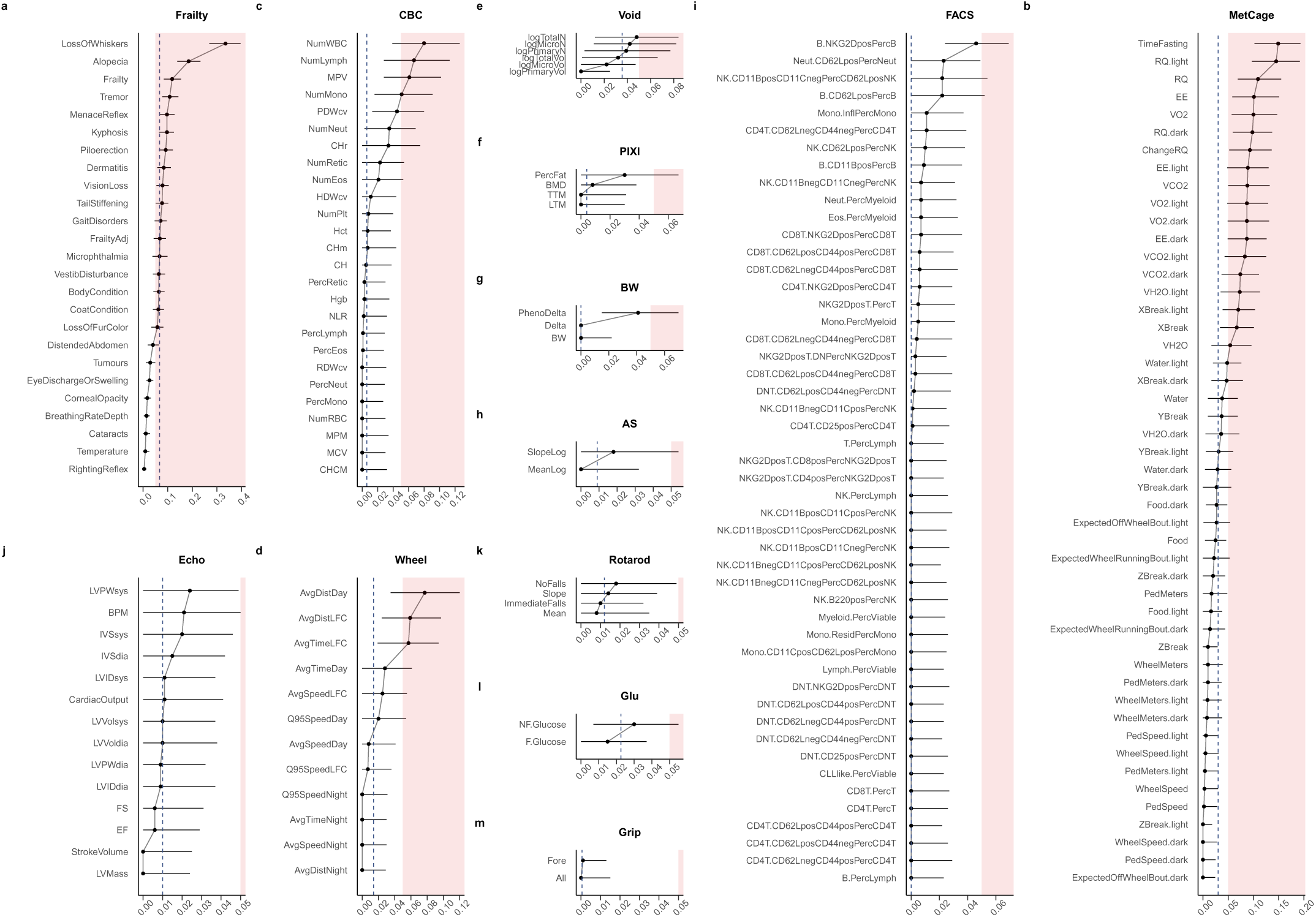
Intra-cage correlations by physiologic trait domain. ICC was estimated via LMM methods applied to longitudinally collected phenotypic trajectories in the DRiDO study. LMM estimated ICC*±*95% confidence interval shown. Dotted vertical line indicates median ICC for the phenotypic domain. LMM specified as diet fixed effect and HID random effect. DRiDO phenotypic trajectories included biannual frailty and body temperature (Frailty; a), annual whole blood analysis (CBC; c), annual metabolic cage (MetCage; b), annual wheel running (Wheel; d), annual bladder function (Void; e), annual body composition (PIXI; f), weekly body weights (BW; g), annual acoustic startle (AS; h), annual immune cell profiling (FACS; i), annual echocardiogram (Echo; j), annual rotarod (Rotarod; k), annual fasting and non-fasting glucose (Glu; l), and annual grip strength (Grip; m). Color indicates co-housing effect values*>*0.05. All quantitative traits excluding bodyweights were corrected for batch effects as described in [22]. LMM estimated ICC and 95% confidence intervals are reported in Supplementary Table 1.

### Impact of co-housing on statistical tests for effects in murine aging studies is modest in practice

Analytic approaches varied for the different studies as appropriate given varying study design complexity, yet none of the primary publications accounted for non-independence by cage. (Lifespan data for two studies have not previously been reported [JAXCC and JAXDO].) Underutilization of analytic methods that account for clustering is not unique to preclinical research (e.g., cancer trials [23]; nutrition and obesity [24]). After incorporating various study design features via pre-analysis batch adjustment or model specification, we found that tests for main effect in each study with and without incorporating a random effects term for housing identifier to capture intra-cage correlation usually showed no practical difference (**Supplementary Table 2**).

Repeated measures designs may wrongly be thought to circumvent statistical artifacts of co-housing by comparing mice to themselves over time. In repeated measures applications, the individual becomes the first level of aggregation and the cage or other clustering unit becomes a second level of aggregation. To arrive at an unbiased estimate of outcome trajectories (how individual units on average change over time) from serial data, the second level of aggregation must be accounted for. For longitudinally collected phenotypic trajectories in the DRiDO study, we found that tests for main effect of diet with and without incorporating a random effects term for housing identifier to capture intra-cage correlation often showed no practical difference (**Supplementary Table 3**). Where tests for main effect of diet conflicted, as was the case for several frailty items, p-value for the main effect of diet was underestimated when co-housing was ignored, likely due to misattribution of variance to diet instead of housing ID, which resulted in underestimated standard errors.

### Sample size guidelines in murine aging research lack broad applicability

Murine lifespan exhibits heterogeneity by strain, sex, and site, as highlighted by previous studies [25–27]. Despite this complexity, researchers may rely on sample size recommendations based on survival data from a single study, inbred strain, or sex ([28–33] cite [34]). Similarly, non-lifespan murine outcomes demonstrate considerable heterogeneity [35–37], yet existing sample size guidelines draw primarily from data sources that are strain-specific (e.g., [38, 39]). Failure to consider important sources of heterogeneity can limit the generalizability of recommendations and may lead to unpowered or inefficient study designs [40, 41]. Comprehensive databases providing sample size recommendations across strains, sexes, and experimental conditions would benefit the field [42], yet even in strain-and sex-matched samples, investigators risk non-negligible effects on statistical validity when implementing existing lookup tables for a simplified version of the study design and extrapolating to the more complex case planned [43]. Simulations demonstrate that applying rules of thumb for murine research of aging that ignore intra-cage clustering may underpower studies and reduce replicability. We simulated 2-arm randomized lifespan studies with null effects and different ICC values. We applied conventional tests for uncensored (LM, LMM, GEE) and censored (COX, COXME) data and calculated empirical power. **Fig. 5** (LM, LMM, GEE) and **Supplementary Fig. 3** (COX, COXME) show how power varies with ICC, model type, and sample size. When generating data from a null fixed effects model, any p*<*0.05 indicates a false positive error. Results showed that tests assuming independence (COX, LM) overestimated power in the presence of positive ICC values. Sample sizes computed by COX and LM models ignoring ICC would be too small for positive ICC, leading to wasted resources and low replicability.

**Figure 5:**
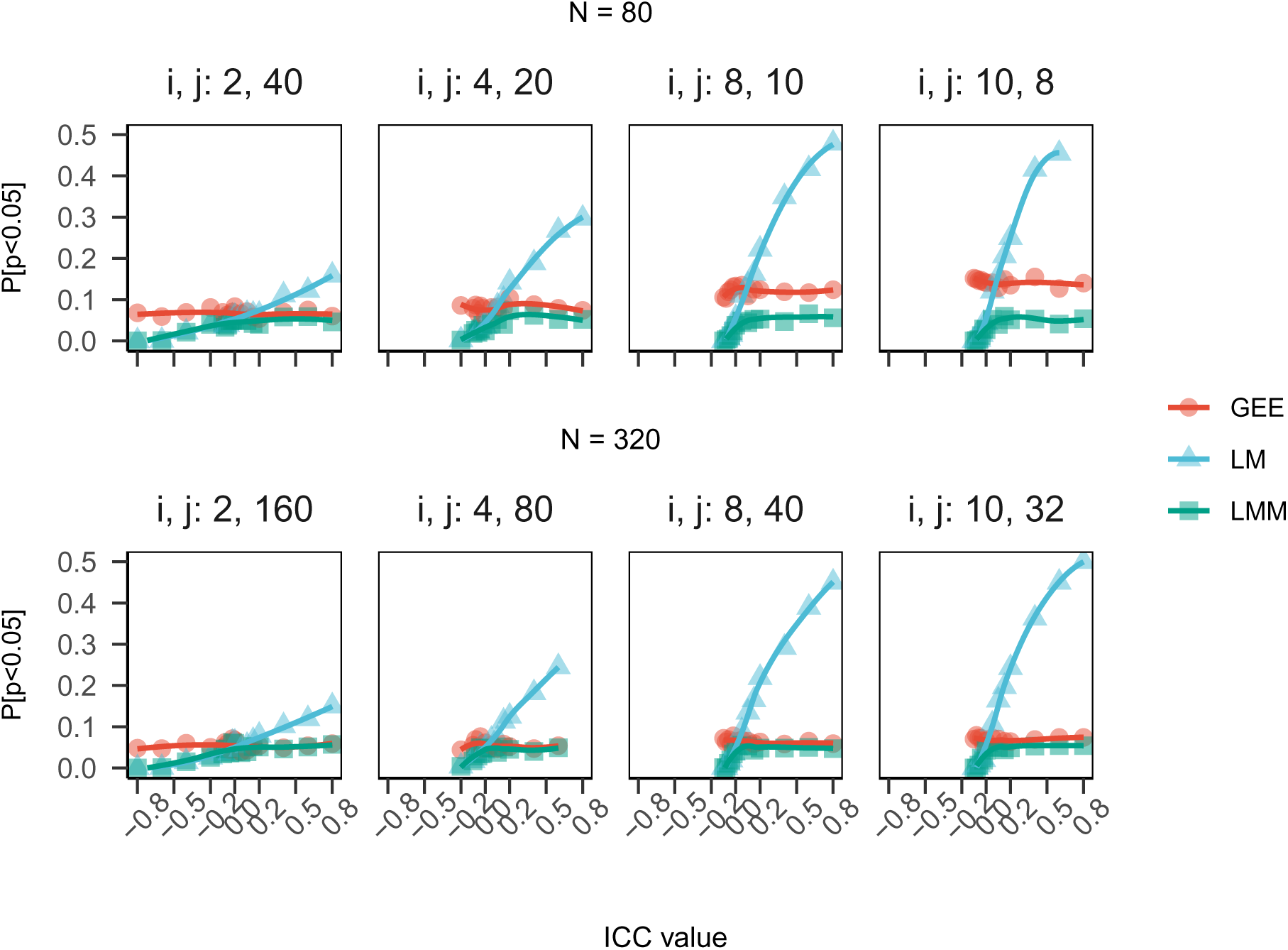
Power curves for simulated trial data with null treatment effects by ICC by model type. Simulations demonstrate that applying rules of thumb for murine research of longevity that ignore intra-cage clustering may under-power studies and reduce replicability. We simulated 2-arm randomized lifespan studies with null effects and different ICC values. We applied conventional tests for uncensored data (LM, LMM, GEE) and calculated empirical power. The figure shows how power varies with ICC, model type, and sample size. When generating data from a null fixed effects model, any p*<*0.05 indicates a false positive error. The results showed that tests assuming independence (LM) over-estimated power in the presence of positive ICC values. Sample sizes computed by the LM models ignoring ICC would be too small for positive ICC, leading to wasted resources and low replicability.

### Outcome dependence results in biased p-value distributions

Why does this overinflation of power occur for LM and for COX models where ICC is non-null? Behavior of the p-value distribution, which is assumed to be uniform from zero to one under the null hypothesis is central to the hypothesis testing paradigm. Via simulation, we show that violated assumptions can cause the p-value distribution to be skewed under the null. For example, if a test assumes ICC=0, as in LM or COX, but ICC=*−*0.1, then the p-value distribution will be skewed towards 1 (**Supplementary Fig. 4**). This will result in a loss of power and an increase in type II errors (failing to reject the null hypothesis when it is false). Conversely, if again a test assumes ICC=0 but ICC=+0.1, the p-value distribution will be skewed towards zero (**Supplementary Fig. 5**). This will result in an overestimate of power and increase in type I errors (rejecting the null when it is true). Therefore, in murine aging studies, it is important to check the assumptions of the test and use appropriate methods to account for any violations. Statistical biases resulting from intra-cage correlations with similar magnitudes as observed in these case studies have important implications for power and reproducibility.

While lifespan outcome is characterized by a single quantitative variable, quantification of health effects or molecular changes in response to aging interventions is often a multi-outcome endeavor (e.g., [22, 44]). With the advent of digital cages capable of 24/7 automated data capture, multi-outcome preclinical aging research is likely to become only more highly dimensional in the near future [45]. It is recommended practice to employ false discovery rate (FDR) adjustment to account for multiple testing. However, when conducting many tests and applying an FDR adjustment, skew in the null p-value distribution can affect the estimation of the proportion of true null hypotheses and the calculation of adjusted p-values [46, 47]. Biased p-value distribution can thus result in either too many or too few false discoveries when applying an FDR adjustment to large-scale simultaneous hypothesis tests. Consequently the importance of considering biased p-value distribution due to non-independence by housing unit under the null hypothesis may be even greater for biomarkers of aging than for lifespan, despite similar observed ICC magnitude.

Sample size guidelines used in murine aging research may be derived from formulas that ignore potential correlation induced by co-housing (e.g., [34]). Quantified ICC in lifespan data pooled across multiple sites, sexes, diets and in hundreds of traits in a large sample of genetically diverse mice allows for improved estimates. For census observed data, assuming the same mean and variance reported in [34] (mean days=912, variance=143^2^) and assuming n-per-cluster is 4 (mean in database=4.07), we estimated n per group as follows: n=39 for ICC=0 (as in [34]; rounded to the nearest 10), n=40 for ICC=0.01, n=45 for ICC=0.05, and n=51 for ICC=0.1. We also provide a small grid of required number of mice per group according to the closed form solution [48] for a 10% effect size by ICC and cluster size for inbred and outbred mice (**Table 3**, **Fig. 6**). Our expectation is that these recommendations will be continuously updated by statisticians in the preclinical aging research community as new data become available. These results imply that strain and sex show large effects on n per group relative to plausible ICC and cluster size for murine aging studies.

**Figure 6:**
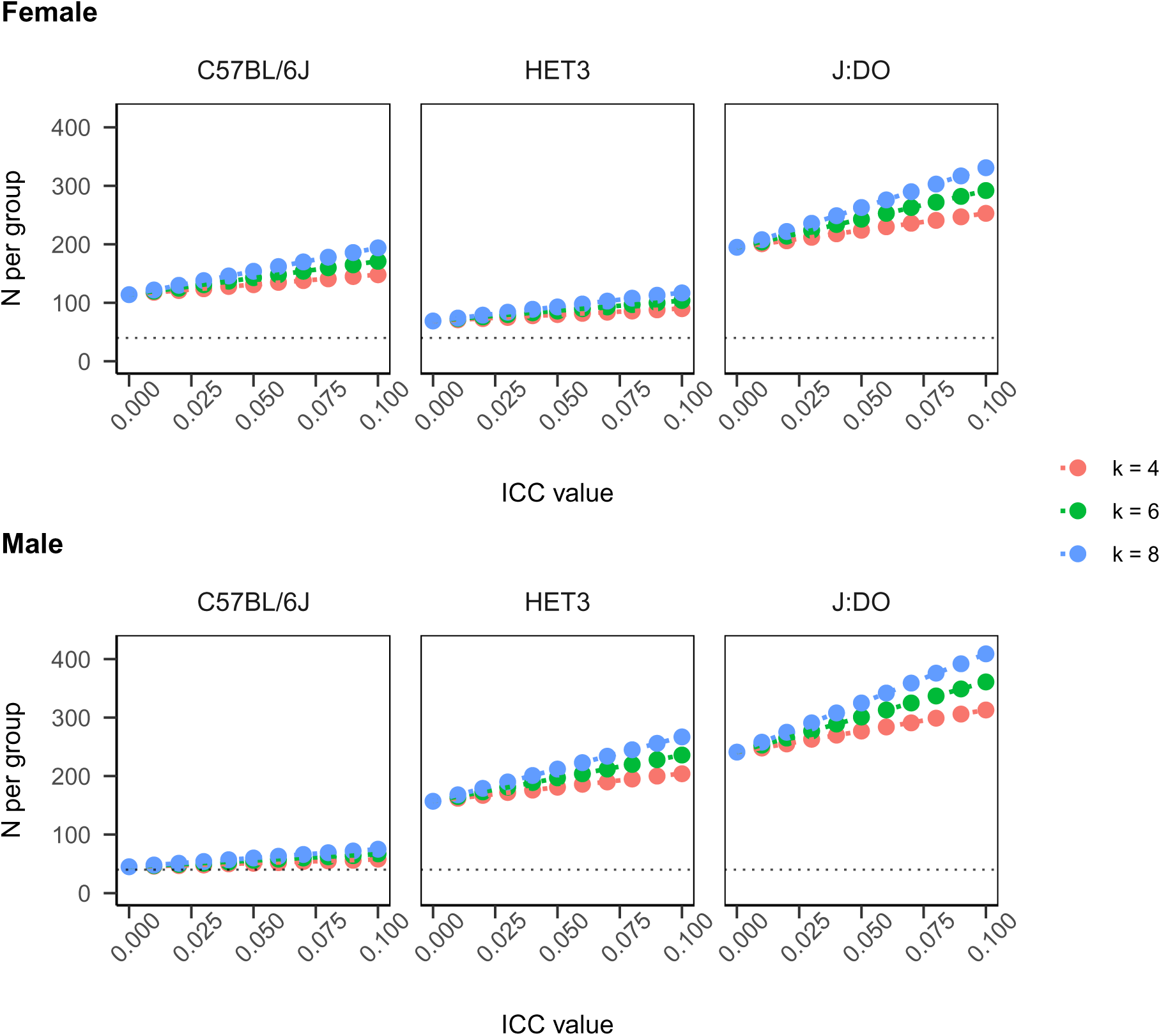
Sample size to detect fractional change in mean outcome under CRT design. The impact of the ICC on the planned trial size is often discussed as being dependent on its magnitude and on the number of subjects recruited per cluster, n, through the so-called design effect (DE), (1) DE = [1 + (n-1)*α*] [48]. For a simple comparison of means in a two-arm trial with equal allocation per group, the DE is used as an inflation factor multiplied by the total sample size for independent data. The table shows a grid of estimates for required number of mice per group according to closed form solution in equation (1) for 10% ES [0.1*×*strain-specific-mean lifespan in database] with census follow-up, and anticipated power 1-*β* of 0.8. Values presented separately by sex, ICC, and cluster size. ICC varied *∈{*0.0, 0.01, 0.05, 0.1*}*, cluster size varied *∈{*4, 6, 8*}*. ICC=intra-class correlation coefficient, ES=effect size, k=cluster size. Results imply that strain and sex show large effects on n per group relative to plausible ICC and k for murine aging studies. Note that source data for mean and variance in C57CL/6 were more sparse than for outbred mice and so are less reliable. Our expectation is that these recommendations will be continuously updated by statisticians in the preclinical aging research community as new data become available.

**Table 3:** Sample size to detect fractional change in mean outcome under CRT design. The impact of the ICC on the planned trial size is often discussed as being dependent on its magnitude and on the number of subjects recruited per cluster, n, through the so-called design effect (DE), (1) DE = [1 + (n-1)*α*] [48]. For a simple comparison of means in a two-arm trial with equal allocation per group, the DE is used as an inflation factor multiplied by the total sample size for independent data. The table shows a grid of estimates for required number of mice per group according to closed form solution in equation (1) for 10% ES [0.1*×*strain-specific-mean lifespan in database] with census follow-up, and anticipated power 1-*β* of 0.8. Values presented separately by sex, ICC, and cluster size. ICC varied *∈{*0.0, 0.01, 0.05, 0.1*}*, cluster size varied *∈{*4, 6, 8*}*. ICC=intra-class correlation coefficient, ES=effect size, k=cluster size. Results imply that strain and sex show large effects on n per group relative to plausible ICC and k for murine aging studies. Note that source data for mean and variance in C57CL/6 were more sparse than for outbred mice and so are less reliable. Our expectation is that these recommendations will be continuously updated by statisticians in the preclinical aging research community as new data become available.

|  | ICC=0 | ICC=0.01 | ICC=0.05 | ICC=0.1 |
| --- | --- | --- | --- | --- |
| **J:DO** |  |  |  |  |
| ES=10%, k=4 | $F = 195, M = 241$ | $F = 201, M = 248$ | $F = 224, M = 277$ | $F = 253, M = 313$ |
| ES=10%, k=6 | $F = 195, M = 241$ | $F = 204, M = 253$ | $F = 243, M = 301$ | $F = 292, M = 361$ |
| ES=10%, k=8 | $F = 195, M = 241$ | $F = 208, M = 258$ | $F = 263, M = 325$ | $F = 331, M = 409$ |
| **HET3** |  |  |  |  |
| ES=10%, k=4 | $F = 69, M = 157$ | $F = 71, M = 162$ | $F = 80, M = 181$ | $F = 90, M = 204$ |
| ES=10%, k=6 | $F = 69, M = 157$ | $F = 73, M = 165$ | $F = 86, M = 197$ | $F = 104, M = 236$ |
| ES=10%, k=8 | $F = 69, M = 157$ | $F = 74, M = 168$ | $F = 93, M = 212$ | $F = 117, M = 267$ |
| **C57BL/6** |  |  |  |  |
| ES=10%, k=4 | $F = 114, M = 45$ | $F = 118, M = 46$ | $F = 131, M = 51$ | $F = 148, M = 58$ |
| ES=10%, k=6 | $F = 114, M = 45$ | $F = 120, M = 47$ | $F = 143, M = 56$ | $F = 171, M = 67$ |
| ES=10%, k=8 | $F = 114, M = 45$ | $F = 122, M = 48$ | $F = 154, M = 60$ | $F = 194, M = 75$ |

## Discussion

In this study, we performed a secondary analysis of mouse lifespan data from six large cohorts. The assembled database comprises over 20,000 observations with lifespan and cage assignment data. These studies include mice of both sexes representing 42 inbred strains and two outbred populations. The studies were carried out in different facilities, at different times and with a variety of housing configurations. Data were gathered from the Mouse Phenome Database (MPD), a public repository of phenotypic data from various mouse strains and experimental settings [49], the Intervention Testing Program, a multi-institutional program to evaluate putative anti-aging therapies [50], as well as our own research in the basic biology of aging as part of the Nathan Shock Centers program [51].

An important problem in any analysis of research data is the potential for bias due to omitted covariates. Isogenic cohorts show considerable phenotypic variation among littermates, despite food, temperature, and husbandry standardization [38]. For practical and scientific reasons mice are often co-housed and as result, individual mice may no longer be independent of each other because they are exposed to common influences besides treatment. These dependencies lead to the unit of analysis problem. In a hypothetical scenario, imagine that mice in one cage receive a specific treatment, while mice in a separate cage serve as controls. Treatment condition becomes entirely confounded with the cage variable. Consequently, it becomes challenging to discern whether observed outcome differences are attributable to the treatment itself or simply variations between cages (temperature, light exposure, etc.).

Observations on individual units within the aggregate may be independent of one another, but we cannot assess this for most murine aging studies because housing identifiers are not published. Standardization of lifespan data reporting would support re-analysis efforts [42]. In this report we used experimental data from six preclinical aging cohorts with available housing identifiers to illustrate the impact (or lack thereof) of statistical dependence of outcome stemming from co-housing. By quantifying intra-cage correlation in murine lifespan studies, we aimed to provide a necessary tool for researchers in the planning phase of future lifespan intervention trials in aging research. As these studies typically have many housing units with small numbers of animals per unit, the impact on analysis outcomes were negligible. Nonetheless there are potential risks to ignoring ICC, including over estimation of power in study design and increased rates of type I error in analysis.

Pooled estimates in this large database are consistent with the general guidance for simulating clustered data in power analyses with ICC=0.05 may be generally appropriate to murine models of aging. LMMs and GEEs are based on asymptotic theory and, therefore, assume many clusters. Failure to meet this assumption results in increased type I error rate. The practicalities of animal husbandry have encouraged good cRCT design in murine aging studies with lifespan outcome (many clusters/cages, small n per cluster/cage), resulting in studies that are largely robust to confounding by clustering. Smaller studies with fewer housing units will be more susceptible to effects of clustering.

We also show, via differences in estimated ICC by study, that this ‘guesstimate’ can be improved upon, especially in outbred mice for which we had very large N. Using a large database encompassing broad surveys of genetic variation, anti-aging therapeutics, and both sexes, we identified that ICC estimates are nonconstant, ranging from negative to null to neutral. The ICC is typically described as ranging from 0 for complete independence of observations to 1 for complete dependence under the assumption that clustering produces similarity or positive correlation among cases within a cluster (the most common presentation in clustered data). While it is true that a negative variance can be due to a mis-specified model, it is also possible that an experimenter may create clusters in which individual units are more dissimilar from one another than might be expected by chance alone, e.g. [52]. A readily apparent example is when individuals compete in a cluster for scarce resources. In this instance, ICC can be negative and lead to an actual Type I error rate lower than the nominal Type I error rate [53]. The two dietary treatment studies allowed us to quantify ICC in exactly this scenario. We demonstrated how researchers might inspect their data for a dispersion effect (negative ICC) via GEE estimation. We show that that GEE is preferable for anticorrelated data based on null distribution of p-values. Proper experimental design depends on predetermination of study size to ensure that hypothesized effects will be detectable. Having estimated ICC for a wide variety of murine aging studies, we carried out simulation studies using housing configurations and levels of ICC that are typical for murine aging studies. We examined implications for power and reproducibility and compare analysis outcomes using methods that do or do not account for ICC. Simulation results suggest that oversimplified estimation procedures that ignore ICC can lead to inefficient designs and unreliable results. Power analyses that assume independent data can provide overly optimistic (too small) estimates of sample size. Pooled estimates in this large database are consistent with the general guidance for simulating clustered data in power analyses with ICC=0.05 may be generally appropriate to murine models of aging. Incorporating ICC supports better-informed power estimation and ultimately experimental design in aging research that is more robust to confounding by intra-cage correlation.

In practice, we have shown that ICC will be small or negligible in murine aging studies of lifespan. Reassuringly, sample size guidelines for this data type that ignore clustering by cage are likely to be only slightly underpowered. We caution that other outcomes were also found to be subject the effects of cohousing. In future work we intend to apply these methods to molecular aging data. Access to more preclinical aging studies that include housing information will contribute to improved design guidelines for future studies, maximizing the efficiency of preclinical aging research by preventing underpowered studies.

## Materials and Methods

### Datasets from lifespan studies

We used data from five previous lifespan studies conducted with support from the National Institute on Aging’s Nathan Shock Center for Excellence in the Basic Biology of Aging and Calico Life Sciences LLC. In all studies, natural lifespan data served as the primary outcome. These data are distinctive in that unique housing unit identifiers were available along with lifespan data; publishing housing identifiers is not standard in the field of pre-clinical aging research. All studies are briefly described in **Table 1**; JAXDO and JAXCC have not been published previously. The aggregated database included:

> The Jackson Laboratory Collaborative Cross Lifespan Study (JAXCC),

> The Intervention Testing Program (JAXITP and UTITP),

> The Jackson Laboratory 32 Strain Lifespan Survey (JAX32),

> The Jackson Laboratory Diversity Outbred Lifespan Study (JAXDO), and

> The Dietary Restriction in Diversity Outbred mice Study (DRiDO).

### Data Extraction

Mouse ID, housing ID, strain or genetic background, sex, diet, treatment group, death status (censored or observed), and lifespan (days) were extracted from each dataset. Variables that might indicate batch effects (group, cohort, generation) were also extracted where available. For data extraction of mortality, negative lifespans were removed (n=6 in JAX32). One strain, Pohn/DeH, was excluded from the JAX32 database to maintain consistency with the anchor manuscript associated with primary data.

Housing ID and Mouse ID were available in the primary dataset for all studies except the ITP. Housing identifiers are not routinely collected as part of the ITP database; these were sought from each site separately and were made available for two of the three ITP sites (JAX and UT). Housing ID files were merged with the ITP lifespan database using unique mouse identifiers. All 9,425 lifespan observations associated with JAX were matched to a housing identifier. There were 9,075 observations in the UM database associated with UT through C2017 and UT provided 8,545 housing identifiers through C2016. 951 observations associated with UT from C2017 and 487 observations associated from UT from C2004-C2006 were excluded due to no available housing identifier, resulting in 7,637 observations associated with UT with raw survival data provided by UM and housing data provided by UT. All lifespans were divided by 30.4 to convert days to months of life. None of these data were conditioned on early survival (e.g., not filtered on survival to 6 months), which may differ from data presentation in a primary publication. Overall, 22,385 mice were included in the final database. Less than 4% of mice were censored; these mice were excluded from analyses.

### Batch correction

To prevent potential biases in interpretation and increase reliability of natural lifespan measurements, we corrected values for batch effects. To quantify batch effects, we fit linear mixed models conditioning on housing unit, strain (for JAX32 and JAXCC), phenotyping or randomization group (for DRiDO, JAXCC, JAXITP, UTITP), sex (all except DRiDO), and batch. These models were estimated in data with far outliers removed to avoid over-adjustment for extreme values [54]. To remove batch effects, we adjusted continuous lifespan outcome through subtraction of batch model coefficients. Batch was defined by the following design factors: by group in JAX32, by generation and cohort in JAXCC, by generation in JAXDO, by generation in DRiDO, and by cohort in ITP [estimated separately for JAX and ITP cohorts]). For DRiDO and JAXCC we determined a reduced model excluding generation was sufficient to control for the batch effects. Among standard statistical packages in R, GEE implementations can only handle one level of clustering at a time. Thus for JAX32 and JAXCC, strain random effects were also removed to maintain a 2-level hierarchical data structure across all lifespan outcomes. Negative adjusted survival values generated by these study-specific models were removed (n*<*10).

### Statistical analysis of lifespan database

Descriptive statistics and boxplots were used to describe housing densities by study and interquartile range of residual survival after adjustment for study factors. Boxplots were used to demonstrate variability in lifespan by study, depicting outcome distributions across randomization group, strain, and sex (n=120 strata).

We applied LMM and GEE methods to each of the studies to estimate ICC in lifespan data. Confidence intervals around LMM ICC estimates were quantified via model-based parametric bootstrap methods. Permutation testing quantified significance of the observed ICC statistic estimated by GEE methods. LMM condition on random effects, while GEE integrates out random effects, a distinction with important downstream effects on analytic results [20]. In the presence of negative intra-cage correlation, LMM will constrain the random effect variance to zero, thereby forcing the statistical model to assume the data are independent and identically distributed (iid), an assumption which is unlikely to hold true. Unlike their conditional counterpart, GEE models work from a marginal framework where interdependency between cages is not implied by a random-effect structure but is explicitly modeled in a covariance matrix. Consequently, GEE is an unbiased estimator of ICC in that it can estimate positive, null, or negative ICC values while LMM is a biased ICC estimator in that it can estimate only positive or null ICC values. We also use permutation testing to visual demonstrate and compare observed versus permuted (null) ICC distribution for each study as estimated by GEE methods. For each study, 1k permutations of the data were generated. P-value plots by study showed values were stable at permutation n*>∼*300.

LMM specifications for studies in the combined database were as follows: For DRiDO, we adjusted outcome for generation batch effects and fit the linear mixed model *y ∼ diet* + (1*|hid*), where *diet* indicates one of five diet protocols and *hid* indicates housing identifier. For JAX32, we adjusted outcome for strain effects and fit the linear mixed model *y ∼ sex* + (1*|hid*). For JAXCC, we adjusted outcome for strain and generation batch effects and fit the linear mixed model *y ∼ sex* + *diet* + (1*|hid*), where *diet* indicates one of two diet protocols. For JAXDO, we adjusted outcome for generation batch effects and fit the the linear mixed model *y ∼ sex* + (1*|hid*). For JAXITP and UTITP studies, we adjusted outcome for cohort year prior and fit the linear mixed model *y ∼ sex* + *rx* + (1*|hid*), where *rx* is a binary variable indicating treatment or control assignment. For the combined model, sex, diet, rx, and study served as fixed effect terms and housing identifier as a random effect term; survival in months was pre-adjusted within study strata as above. GEE specifications: Same as above.

After incorporating various study design features via pre-analysis batch adjustment and model specification, we tested for main effect study factors specific to each experimental design with and without incorporating a random effects term for housing identifier to detect bias in statistical tests for lifespan that do not account for co-housing. Significance of categorical predictors sex, strain, diet, and/or treatment (as relevant for each study, see above) was assessed via F-tests (type III sum of squares) for linear models with and without a housing identifier random intercept using the “joint tests” function in the emmeans R package ([55]).

### Statistical analysis of DRiDO phenotypes

Data for the DRiDO study, a comprehensive investigation of dietary restriction interventions in Diversity Outbred mice, were available via a public data repository in keeping with Open Science principles [56]. Quantitative traits other than bodyweights were corrected for batch effects as described in Di Francesco et al. [22]; see this paper for further details about phenotyping protocols and experimental design. We applied LMM methods to all physiological measurements in DRiDO, resulting in ICC estimates across hundreds of physiologic measurements of aging in a representative (i.e., genetically diverse) dataset. Because phenotypes were collected longitudinally across the lifespan, these data types could by hierarchically clustered both by housing identifier and by animal (**Fig. 1**). Among standard statistical packages in R, GEE implementations can only handle one level of clustering at a time, thus ICC was only estimated via LMM for the longitudinal analysis of DRiDO phenotypes. LMMs were specified with additive fixed effects for diet and age and two random intercepts, housing identifier and animal identifier. ICC specific to the housing identifier random effect for the LMM model with both clustering levels was estimated via the proportion of variance method, which has a lower bound of 0. As for lifespan outcome, confidence intervals around LMM ICC estimates were quantified via model-based parametric bootstrap methods and significance of main effect study factor (here, diet) was assessed via F-tests. These LMMs and post-estimation contrasts were re-run without the housing identifier random intercept to discover any instances where scientific interpretation would be affected by ignoring co-housing.

### Closed Form Estimation of Sample Size

Impact of the ICC on planned trial size is often discussed as being dependent on its magnitude and on the number of subjects recruited per cluster, n, through the so-called design effect (DE), (1) *DE* = [1 + (*n −* 1)*α*] [48]. For a simple comparison of means in a two-arm trial with equal allocation per group, the DE is used as an inflation factor multiplied by the total sample size for independent data. We derived a grid of estimates for required number of mice per group according to closed form solution in equation (1) for 10% effect size (ES) with census follow-up, and anticipated power 1-*β* of 0.8. ES was specified as 0.1*×*strain-specific mean in database. Values were estimated separately by sex, ICC, and cluster size. ICC varied *∈{*0.0, 0.01, 0.05, 0.1*}*, cluster size varied *∈{*4, 6, 8*}*. Due to the controlled environment of preclinical aging research, longevity data may be census observed or nearly so. For a discussion of sample size estimation in clustered survival data with censoring, see **Supplementary Note**.

### Simulation Study

A Monte Carlo simulation study was used to evaluate the statistical appropriateness of various analytic approaches for plausible two-arm randomized preclinical lifespan studies. Plausibility was determined by concordance with the published studies included in our database, described above. The aim was to demonstrate estimation errors that are made in statistical analysis when data are treated as independent, ignoring clustering inherent to the study design of lifespan studies in aged mice. Accuracy of regression parameter estimates, coverage of 95% CIs, and frequency of Type I and Type II errors were computed for each model type. For detailed methods, see **Supplementary Methods**.

## Acknowledgments

This work was supported by the National Institutes of Health (Nathan Shock Centers of Excellence in the Basic Biology of Aging program, grant number AG38070 to G.C.) and Calico Life Sciences LLC (Dietary Intervention of Aging in Genetically Diverse Mice, sponsored research funding number CALICO-GAC-06 to G.C.). We acknowledge the JAX Nathan Shock Center Animal and Phenotyping Core team and the JAX Interventions Testing Program team for their expertise in animal husbandry, data collection, and data curation, and Matthew Vincent for assistance with high performance computing. We thank Martin Mullis and Cynthia Kenyon of Calico Life Sciences LLC for providing helpful comments on an earlier draft of the manuscript. Additionally, we acknowledge Dr. Richard Miller of the University of Michigan for providing access to ITP survival data and Dr. James Nelson of the San Antonio Nathan Shock Center for sharing supplementary ITP housing unit identifier data.

## Author Contributions

GC: conceptualization, funding acquisition, data acquisition, project administration, methodology, formal analysis, writing. AL: methodology, formal analysis, writing.

## Data Availability Statement

All analyses were performed using the R statistical programming language [57]. Lifespan data and code used to generate tables, figures, and reported results can be found on figshare (DOI: 10.6084/m9.figshare.26485633). Data for DRiDO phenotypes can be found on the online data repository for the anchor manuscript [56].

