## Supporting Information for "Quantifying the Impact of Co-Housing on Murine Aging Studies"

Supplementary Table 1

Supplementary Table 2

Supplementary Figure 1

Supplementary Figure 2

Supplementary Figure 3

Supplementary Figure 4

Supplementary Figure 5

Supplementary Methods

Supplementary Note

### Extended Data

**Supplementary Figure 1:** Distribution of residual lifespan by housing unit in JAXDO after after accounting for all other study design effects. Each boxplot represents one housing unit, displaying the median, interquartile range (25th to 75th percentile; IQR), and 1.5$\times$ IQR. Boxplots of lifespan by housing ID were sorted by median residual lifespan (highlighted in red). Taller boxplots within study indicate cages with larger intra-cage variability and non-zero slope of the medians indicates intra-cage correlation within the study.

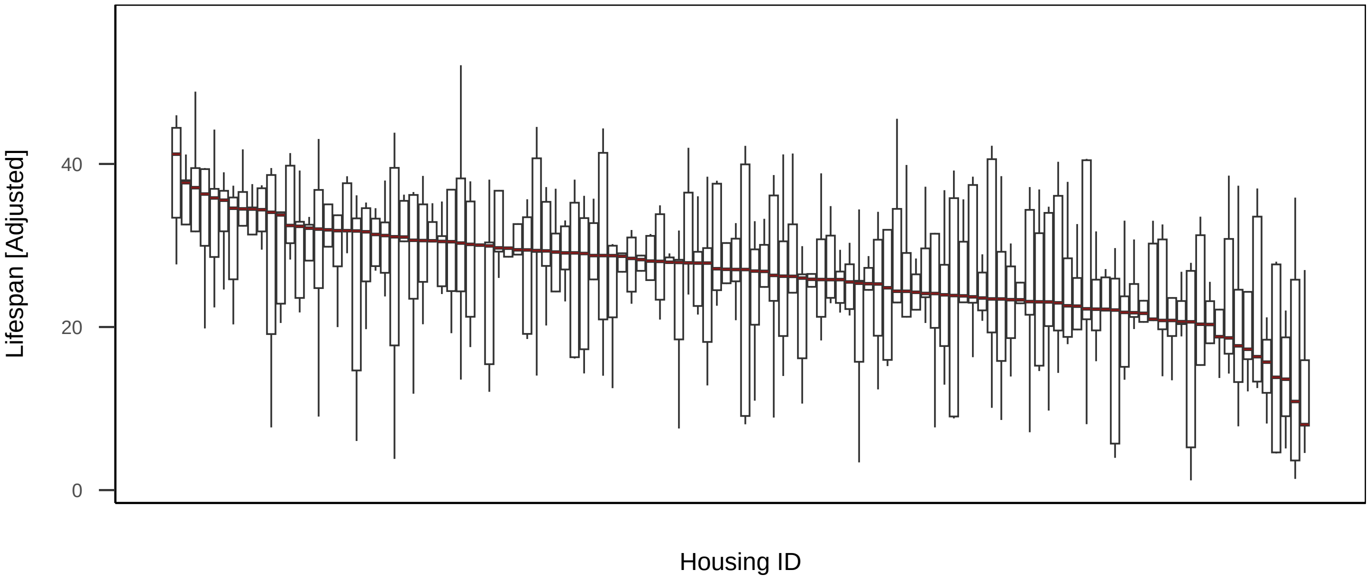

**Supplementary Figure 2:** Permutation tests for non-independence by cage for lifespan outcome across included cohorts. For each cohort, a set of 1000 permuted datasets was generated by randomly assigning housing unit identifiers. GEE models were fit to each permutated dataset and to the observed data to estimate ICC for null and observed distributions. Comparing values for the observed ICC to the distribution of permutated ICCs indicated statistical significance.

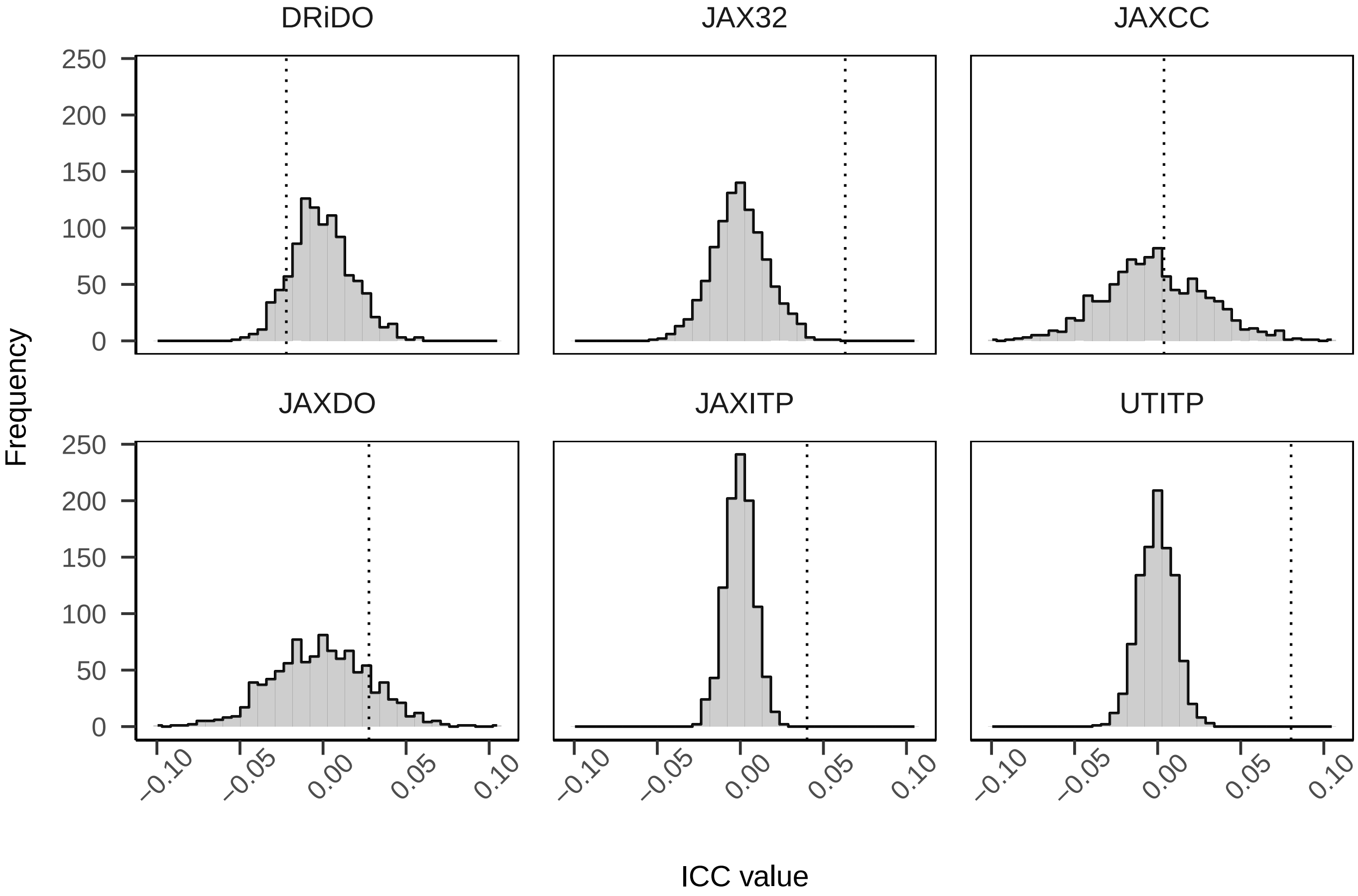

**Supplementary Figure 3:** Power curves for simulated trial data with null treatment effects by ICC by model type. Simulations demonstrate that applying rules of thumb for murine research of longevity that ignore intra-cage clustering may underpower studies and reduce replicability. We simulated 2-arm randomized lifespan studies with null effects and different ICC values. We applied conventional tests for censored data (COX, COXME) and calculated empirical power. The figure shows how power varies with ICC, model type, and sample size. When generating data from a null fixed effects model, any p¡0.05 indicates a false positive error. The results showed that tests assuming independence (COX) overestimated power in the presence of positive ICC values. Sample sizes computed by the COX models ignoring ICC would be too small for positive ICC, leading to wasted resources and low replicability.

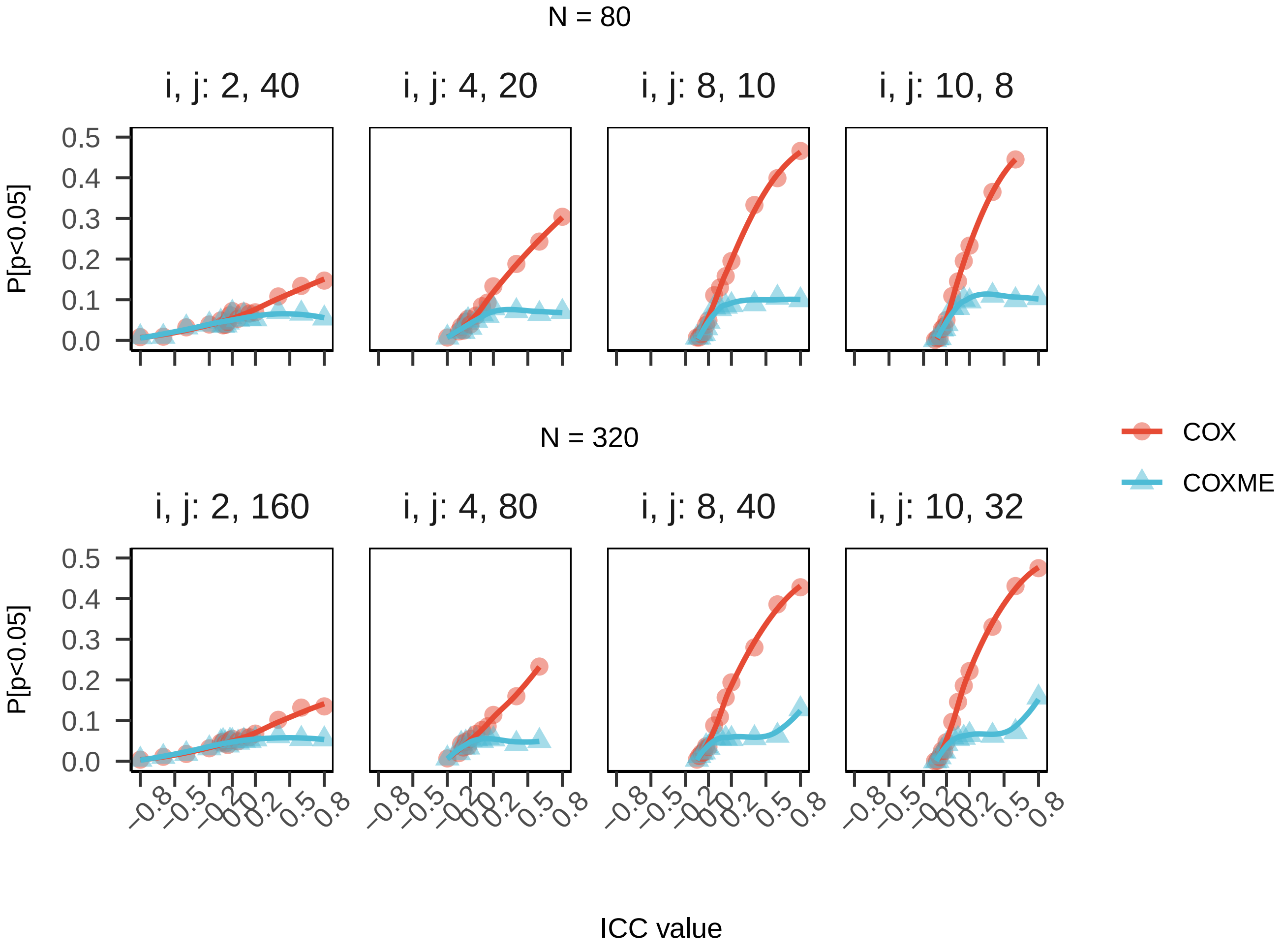

**Supplementary Figure 4:** P-value histograms from 1000 simulations of randomized data with null treatment effect and negative ICC. The distribution of the p-value is a measure of how likely it is to observe a certain p-value or lower under different scenarios. Under the null hypothesis, the p-value has a uniform distribution over [0, 1], meaning that any p-value is equally likely to occur. Under the alternative hypothesis, the p-value has a skewed distribution that depends on the true value of the parameter being tested and the sample size. The more skewed the distribution is towards 1, the lower the power of the test, because large p-values are more likely to occur when the null hypothesis is false. Notes: COX= Cox Proportional Hazards Model; COXME = mixed effects COX; GEE = generalized estimating equations; LM = linear model; LMM = linear mixed model, n.per = number per cluster; negative ICC = -0.1.

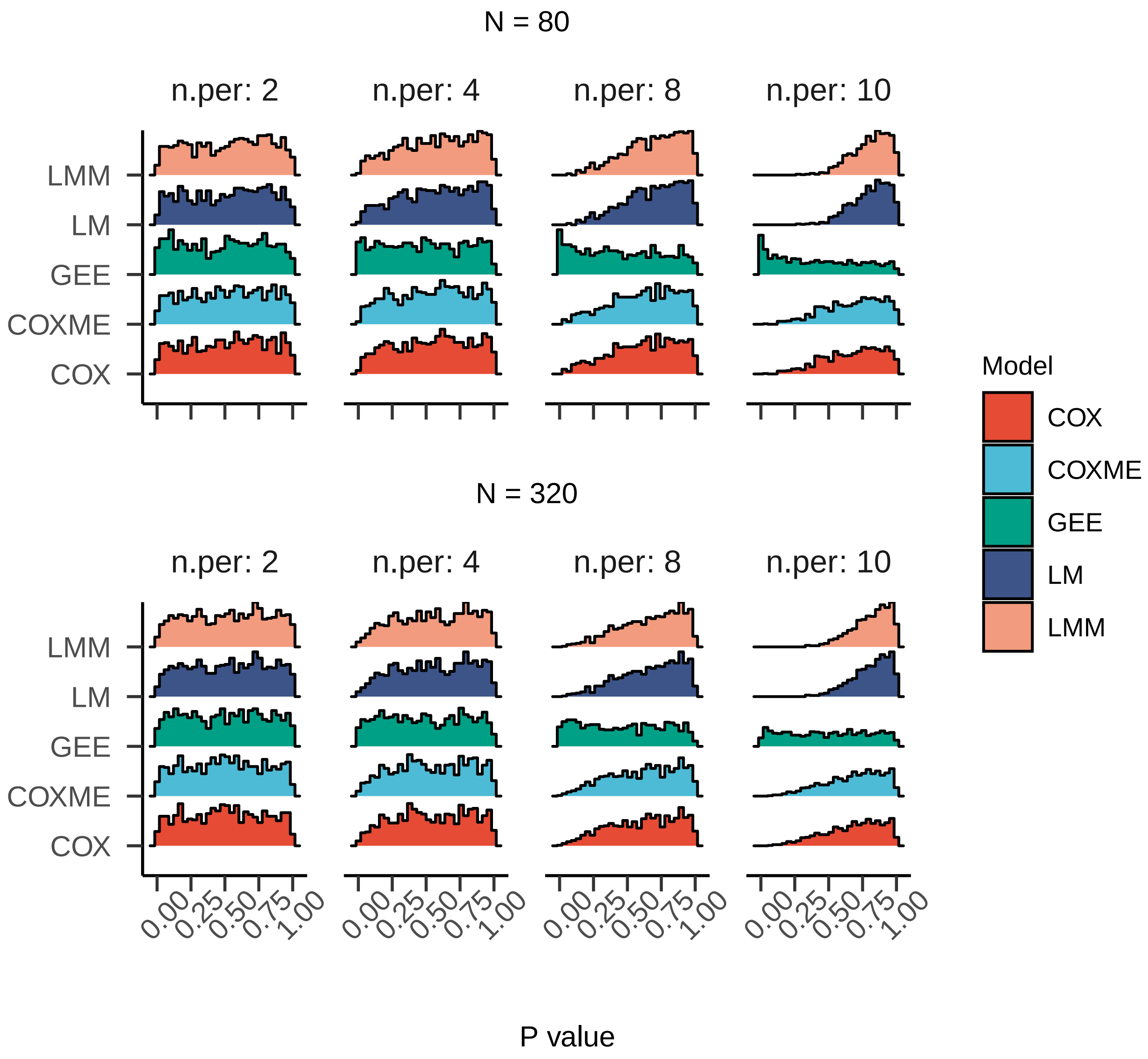

**Supplementary Figure 5:** P-value histograms from 1000 simulations of randomized data with null treatment effect and positive ICC. The distribution of the p-value is a measure of how likely it is to observe a certain p-value or lower under different scenarios. Under the null hypothesis, the p-value has a uniform distribution over [0, 1], meaning that any p-value is equally likely to occur. Under the alternative hypothesis, the p-value has a skewed distribution that depends on the true value of the parameter being tested and the sample size. The more skewed the distribution is towards 0, the higher the power of the test, because small p-values are more likely to occur when the null hypothesis is false. Notes: COX= Cox Proportional Hazards Model; COXME = mixed effects COX; GEE = generalized estimating equations; LM = linear model; LMM = linear mixed model, n.per = number per cluster; positive ICC = +0.1.

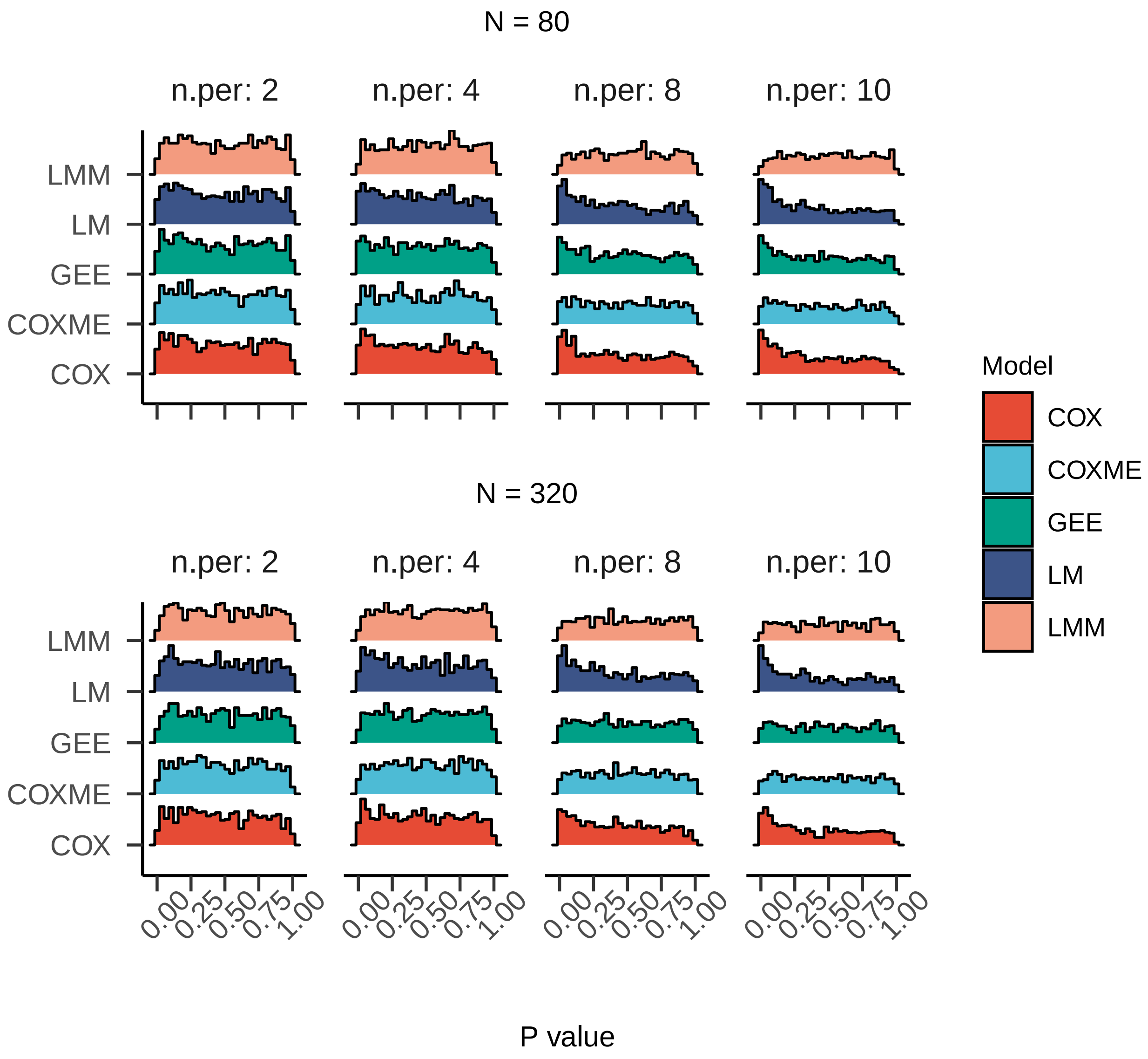

**Supplementary Table 1: Intra-cage correlations by physiologic trait domain in DRiDO.** ICC was estimated via LMM methods applied to longitudinally collected phenotypic trajectories in the DRiDO study. LMM estimated ICC$\pm$95% confidence interval shown. All quantitative traits excluding bodyweights were corrected for batch effects as described in [22]. See DRiDO_all_traits_iccs.csv

**Supplementary Table 2: Type III tests for main effect in each study with and without incorporating a random effects term for housing identifier to capture intra-cage correlation.** After incorporating various study design features via pre-analysis batch adjustment and model specification, we found type III tests for main effect in each study with and without incorporating a random effects term for housing identifier to capture intra-cage correlation usually showed no practical difference. Notes: 1 = singularity; HID = housing identifier, rx = treatment group (control or not control). Estimates may vary from main publications as censored observations were removed and estimates were derived from linear models.

| Study | Batch Adjustment | Fixed Effect(s) | Random Effect | LM p-value | LMM p-value |
|---|---|---|---|---|---|
| DRiDO | generation | diet | HID | <1e-7 | <1e-7[1] |
| JAX32 | cohort, strain | sex | HID | 0.729 | 0.701 |
| JAXCC | generation, cohort, strain | sex, diet | HID | 3.6e-4, 0.005 | 5.5e-4, 0.007 |
| JAXDO | generation | sex | HID | 0.204 | 0.238 |
| JAXITP | cohort | sex, rx | HID | <1e-7 | <1e-7 |
| UTITP | cohort | sex, rx | HID | <1e-7 | <1e-7 |

**Supplementary Table 3: Type III tests for main effect by physiologic trait in DRiDO.** P-values for test for diet across >200 longitudinally collected phenotypes were estimated from linear mixed models with additive fixed effects for age and diet and a random term for mouse identifier with and without incorporating a random effects term for housing identifier (p_diet_w_hid and p_diet_wo_hid, respectively) to capture intra-cage correlation. Estimates may vary from main publications as censored observations were removed and estimates were derived from linear models. All quantitative traits excluding bodyweights were corrected for batch effects as described in [22]. See DRiDO_all_traits_pDiet.csv

### Supplementary Methods

Two total sample sizes were considered: n=80 reflecting the smallest 2-arm randomized trial that would meet published recommendations for lifespan studies in aged mice (Liang, 2003) and n=360 reflecting the largest n-per-group observed within a site in the database (n=160, DRiDO). Number of mice per cluster (n) was varied with attention to practical limits to cluster size such that n $\in\{2,4,8,10\}$ per cage. Number of clusters (housing units) (N) was varied to assess effect of small and large sample sizes such that for a two-arm randomized study with 80 mice total, N $\in\{8, 10, 20, 40\}$, and for a two-arm randomized study with 320 mice total, N $\in\{32, 40, 80, 160\}$. We simulated the null case such with treatment effect equal to zero. Under this null case, any statistically significant regression estimates for treatment observed would be a misattribution of cage effects to the treatment coefficient. Error variance $\sigma^2$ was fixed to one (i.e., outcome was assumed to be z-scored). Simulated intraclass correlation coefficient (ICC) values included positive values indicating similarity within cage, negative values indicating dissimilarity within cage, and null values indicating cage was uninformative for simulated lifespan outcome, with ICC varied from -0.8 to +0.8. Regression parameters and their p-values for LM, LMM, GEE, Cox, and CoxME were estimated. From these, bias, coverage, and power were estimated. Simulated generation of datasets and model estimation were repeated 1000 times per combination and summarized as means (for estimates of treatment effect, p-values, treatment parameter confidence interval bounds, estimates of ICC and its confidence interval bounds) or proportions (coverage rate, power estimates). Coverage rate was computed as the proportion of data replications where the 95% confidence interval contained the true value parameter. Power estimates were computed as the proportion of replicates with estimated p-values$<$0.05.

To simulate data, we generated compound symmetric covariance matrices within cluster/cage for each ICC. With the multivariate normal distribution specified, samples were obtained using the mvrnorm() function in the MASS package (version 7.3.60; Venables and Ripley, 2002). ICC$\times$n-per-cage combinations resulting in a non-positive definite covariance matrix were excluded by setting a lower limit ICC for each n-per-cage = $-\sigma^2/(n-1)$. Regression parameters and their p-values for LM, LMM, GEE, Cox, and CoxME models were estimated using the lm() function in the stats package (version 4.3.1; R Core Team, 2023), the lmer() function in the lmerTest package (version 3.1.3; Kuznetsova et al., 2017), the geeglm() function in the geepack package (version 1.3.9; Halekoh et al., 2006), the coxph() function in the survival package (version 3.5.5; Therneau, 2023; Therneau and Grambsch, 2000), and the coxme() function in the coxme package (version 2.2.18.1; Therneau, 2022), respectively. Confidence intervals for study design effects were estimated directly from LM, LMM, GEE, and Cox model fits and via the tidy.coxme() helper function in the ehahelper package for CoxME model fits (version 0.3.9999; Junkka, 2020). ICC estimates and their confidence intervals were extracted from GEE fits directly as ICC correlation estimate $\pm$1.96*correlation standard error. ICC estimates and their confidence intervals were computed from LMM models via the icc() function in the performance package (version 0.10.4; Lüdecke et al., 2021), which estimates ICC confidence intervals based on bootstrapped samples. For computational efficiency, simulations were run in parallel using the future_pmap() function in the furrr package (version 0.3.1; Vaughan and Dancho, 2022) on a compute cluster with 32 cores. All analyses were conducted using the R Statistical language (version 4.3.1; R Core Team, 2023).

### Supplementary Note

Published sample size tables are an efficient solution to identify sample size for simple research designs, but we are not aware of a closed form solution appropriate for murine aging lifespan studies that can accommodate the structured (i.e., correlated) data type with censoring. For clustered longevity data with censored observations, sample size estimation methods for models that assume constant hazards as in expansions of Freedman (1982) or Schoenfeld (1981), e.g., Jahn-Eimermacher (2012), and sample size estimation methods for models that assume constant additive effects over time, e.g., Blaha (2021), do not reflect the expected hazard distribution of death across the age time scale. Further, extensions to Schoenfeld rely on forms of ICC which contain only partial information of the correlation structure underlying survival times (Zong and Cook, 2015). Sample size approaches for parametric models that allow for non-constant event rates have been proposed, such as for the Weibull distribution (Heo 1998; Wu 2015; Han 2023), the Gompertz distribution (Han 2023), and the generalized gamma ratio distribution more broadly (Phadnis 2016), but these are rarely implemented if at all (Jachno 2019), and these approaches do not accommodate clustered data. Murine lifespan studies with censored observations, nonconstant baseline hazards across time and/or study conditions, and shared frailty effects by housing unit, will require simulation methods to estimate sample size n cages and n-per-cage.
